# Spatiotemporal Profiling of Seed-Associated Microbes of an Aquatic, Intermediate Recalcitrant Species, *Zizania palustris* L. and the Impact of Anti-Microbial Seed Treatments

**DOI:** 10.1101/2022.04.20.488936

**Authors:** Clare Gietzel, Lillian McGilp, Jennifer A. Kimball

## Abstract

Northern wild rice (NWR; *Zizania palustris* var. *interior*) (NWR) is an ecologically important, annual, aquatic species native to North America, that is also cultivated in irrigated paddies in Minnesota and California. NWR seeds are desiccation-sensitive and must be kept in hydrated conditions, in which viability can decline rapidly within the first two years of storage. During this period, microbial growth is rampant and the relationship between these communities and the viability of NWR seed is unclear. In this study, we cultured and identified bacteria and fungi found in 27 NWR seed stocks, collected over a five-year period from three locations and four NWR genotypes. Results revealed that microbial communities were heavily dependent on seed viability and communities began shifting after one year of seed storage. Fungi became more prominent as years in storage increased suggesting that fungi begin to outcompete bacteria. Spatial analysis of locations and genotypes revealed a core set of microbes found across locations and genotypes. *Penicillium* and *Pseudomonas* were ubiquitous in NWR hydrated seed storage. We also evaluated the efficacy of four antimicrobial treatments at reducing microbial growth in hydrated NWR seed storage. These treatments did not reduce or drastically change microbial growth or seed viability. However, previously undetected microbes were identified after treatments, which suggested a disruption to the major constituents of the microbiome. Overall, this study identified common microbial constituents found in an aquatic, recalcitrant species, adapted for a cold climate, during seed storage and provides a foundation for future studies to evaluate the effect of microbial communities on NWR seed viability during hydrated storage.

## Introduction

Northern wild rice (NWR; *Zizania palustris* var. *interior*) (NWR) is an ecologically important, annual, aquatic species native to North America, that is also cultivated in irrigated paddies in Minnesota and California (Grombacher et al., 1997; Oelke and Porter, 2016; Biesboer, 2019; Porter, 2019). NWR seed is dormant at harvest, like many non-domesticated grasses, and is considered intermediately recalcitrant, or desiccation sensitive, which is fairly unusual for members of the *Poaceae* family (Cardwell et al., 1978; Probert and Longley, 1989; Pence, 1995; Pammenter and Berjak, 1999; Dickie and Pritchard, 2002). Due to its unique seed physiology, NWR is typically stored *ex-situ* in the dark and submerged at near-freezing temperatures, from harvest until planting the following year, to mimic the species’ natural aquatic environment (Simpson, 1966; Hayes et al., 1989; Kovach and Bradford, 1992). In these storage conditions, seed viability and germination rapidly decline, particularly once seed dormancy is broken, ~3-6 months post-harvest (Simpson, 1966; Grombacher et al., 1997; Berjak and Pammenter, 2008a; McGilp et al., 2020). This decline is a major limiting factor in *ex-situ storage*, such as seed banks, as well as for the maintenance of germplasm in research programs (Raven et al., 2013).

One potential cause of the rapid decline of NWR seed viability is the proliferation of microbial growth during submerged storage (Berjak and Pammenter, 2008b; Makhathini, 2017; McGilp et al., 2020). Microbial growth, particularly fungal growth, is a main factor in the deterioration of orthodox seed during storage (Christensen and Kaufmann, 1965; Kirkpatrick and Bazzaz, 1979; Khairnar et al., 2011; Abo EL-Dah et al., 2016). The composition of storage-associated microbes of orthodox seed can be affected by a number of factors including seed moisture content, storage temperature, plant species, and plant-microbe or microbe-microbe interactions (E. Welty, 1987; Nelson, 2004b; Cottyn et al., 2011; Truyens et al., 2015; Adam et al., 2018). However, few studies have evaluated the microbiome of recalcitrant seed stored in hydrated conditions. Mycock (1990) found that over a period of 21 days *Fusarium* was the predominant seed-associated microbe from seven recalcitrant species (Mycock and Berjak, 1990). More recently, Makhathini (2017) conducted a more comprehensive study of the seed microbiomes of three recalcitrant tree and bush species: *Protorhus longifolia*, *Trichilia dregeana*, and *Garcinia livingstonei*, and the efficacy of seed treatments on the reduction of microbial growth. To the best of our knowledge, no such studies, evaluating the effect of the seed microbiome on seed viability have been conducted using recalcitrant seed from aquatic species.

Due to the interaction between the microbiome and the viability of seed, antimicrobial seed treatments have long been of interest to researchers. As far back as 1651, Samuel Hartlib noted that the brining and subsequent liming of cereal grains reduced the prevalence of smut, resulting in a yield increase in the following growing season (Smith and Secoy, 1976; Hartlib, 1651). Today, the use of antimicrobial seed treatments for the improvement of seed and seedling health is widespread, especially as commercial fungicide seed treatments (FST) have become more prevalent (White and Hoppin, 2004; Lamichhane et al., 2020). In corn, for example, essential oils from mustard seed (*Brassica campestris*), black cumin (*Nigella sativa*), and neem (*Azadirachta indica*) have been used to significantly reduce the growth of major fungal plant pathogens in the genera *Aspergillus*, *Fusarium*, *Alternaria*, and *Dreschelera* (Ghafoor & Khan, 1976; Mirza & Qureshi, 1982; Sitara, Niaz, Naseem, & Sultana, 2008). Sodium and calcium hypochlorite have also been used as antimicrobial seed treatments to reduce the pathogenic load that leads to a decrease in seed quality and an increase in human illness (Sauer, 1986; Schultz & Gabrielson, 1986; Stewart, Reineke, Ulaszek, & Tortorello, 2001).

Although common in orthodox seed, there are few studies that address the use of antimicrobial treatments for the storage of recalcitrant seed. Most studies conducted with recalcitrant species, have utilized surface sterilization treatments for the long-term storage of tissue culture, rather than seed (Engelmann, 2012; Farzana, Palkadapala, Meddegoda, Samarajeewa, & Eeswara, 2008; Nower, 2013). A more recent study by Makhatini (2017) showed that some chemical fungicides were effective at reducing the growth of fungal isolates, initially collected from three recalcitrant species. The study also showed that when one fungicide was used as a seed treatment it improved seedling vigor, when used in combination with a surface decontaminant (NaOCl) and seed encapsulation (Makhatini, 2017).

In this study, we calculated the germination and viability of NWR seed stored across 5 years. In addition, the identification of culturable microbes present in NWR seed was evaluated using 16S ribosomal RNA (16S rRNA) and internal transcribed spacer (ITS) sequences. Seed germination and microbial communities were compared across varieties, locations, and years in storage. Antimicrobial treatments were then applied to NWR seed prior to storage and the effect of treatments was assessed through germination as well as the comparison of microbial genera present prior to and following treatments.

## Materials and Methods

### Microbial Profiling

#### Plant Materials

For this experiment 28 seed lots were tested. Seeds from three NWR varieties, Barron, Itasca-C12, and Itasca-C20, and one elite breeding line, FY-C20, were hand harvested across five years, 2014 to 2018, and from three locations, Clearbrook, Waskish, and Grand Rapids, MN (Table S1). The locations chosen include the University of Minnesota, North Central Research and Outreach Center, as well as two of the largest areas for NWR production. After each year’s harvest, seed was processed, placed in plastic zipper top storage bags with distilled H_2_O, and stored at 3 °C in the dark.

#### Microbial Isolation

Three replications consisting of three individual NWR seeds per genotype and location were placed on both nutrient agar (NA) amended with 50 mg/L cycloheximide and on potato dextrose agar (PDA) amended with 300 mg/L of streptomycin. Cycloheximide (Siegel and Sisler, 1963) and streptomycin (Ark, 1954) have fungicidal and bactericidal activity, respectively, and were used to reduce fungal growth on NA and bacterial growth on PDA. Plates were placed in the dark at 23°C and evaluated every 24 hours for signs of new microbial growth including changes in color, texture, consistency, or growth pattern. Fungi emerging from seeds were purified by the transfer of a hyphal tip as previously described (Leyronas et al., 2012) or a single spore (Leyronas et al., 2012). Bacteria were subcultured to obtain single colonies (Stevenson, 2006). To store microbial samples long term, fungal samples were placed in a 50/50 (v/v) solution of 15% skim milk and 20% glycerol, while bacterial samples were placed in a 15% glycerol solution. After sample preparation, all samples were placed in storage at −80°C.

#### DNA Extraction

Fungal isolates were re-grown from freezer stocks on PDA, amended as described above, and DNA was extracted using the Qiagen DNeasy Plant Mini Kit (Qiagen, Valencia, CA, USA). For bacterial isolates, a small amount of bacteria, separate from the samples transferred to long term storage, was placed in 300 μl of Tris-EDTA (TE) buffer (10 mM Tris-HCl, 1 mM disodium EDTA, pH 8.0) to preserve the cells before DNA extractions. Bacterial DNA was extracted using the QIAamp DNA Mini Kit (Qiagen, Valencia, CA, USA). The concentration of all microbial DNA was quantified using a NanoDrop 2000 microvolume spectrophotometer (Thermo Fischer Scientific, Waltham, MA).

#### PCR and Sequencing

The internal transcribed spacer (ITS) was amplified from fungal isolates using the ITS1F (Gardes et al., 1993) and ITS4 (Bruns et al., 1990) primers. For bacterial DNA, 16S ribosomal RNA (16S rRNA) was amplified using the 27F (Lane, 1991) and 1492R (Turner et al., 1999) primers. Each reaction contained 50 ng template DNA, 12.5μL GoTaq Green Master Mix (Promega Corporation #M7123, Madison, WI), 10 μM forward and reverse primers, and nuclease free water. Amplifications were run on a T100™ Thermal Cycler (Bio-Rad, Hercules, CA). Thermocycler conditions for ITS primers were as follows: a 5 min denaturation step at 95°C; 34 cycles of 95°C for 30s; 55°C for 30s; 72°C for 60s; and a final extension step of 72°C for 10 min. Conditions for 16S rRNA primers included the following: a 5 min at 95°C; 30 cycles of 95°C for 30s; 58°C for 30s; 72°C for 50s; and a 10 min extension step at 72°C. Amplicons were sent to Molecular Cloning Laboratories in San Francisco, California for PCR clean up using ExoSAP-IT™ enzymatic PCR clean-up (Thermo Fischer Scientific, Waltham, MA) and Sanger sequencing on a ABI 3730X DNA analyzer (Thermo Fischer Scientific, Waltham, MA).

#### Germination and Tetrazolium Tests

Seed viability was estimated using both germination and tetrazolium tests. For germination tests, ~20-30 seeds were rinsed in distilled H_2_O, placed in a new Ziploc bag, and positioned in a growth chamber for 3 weeks with 15 hr of light at 18°C and 9 hr of darkness at 9°C. Germination was defined as coleoptile emergence of 1 cm. The tetrazolium tests were conducted according to the Association of Official Seed Analysts 2000 - Tetrazolium testing handbook protocol (Association of Official Seed Analysts, 2000). Seeds that did not germinate during germination testing were bisected down the long axis of the seed to expose the embryo and suspended in a 0.1% solution of tetrazolium blue chloride (Sigma-Aldrich #88190, St. Louis, MO) for 2 hrs. Seeds were then rinsed and observed for staining of the embryo, which indicated metabolically active tissue that could potentially germinate into a viable plant.

#### Data Analysis

To compare germination between years, genotypes, and locations, averages and standard errors were calculated for each year, genotype, and location using R Studio version 1.2.5001 (R Studio Team, 2021). Tetrazolium test results were treated as a binary; as long as at least one seed per sample stained, it was considered to be a viable seed set. Geneious Prime was used to trim 16S and ITS DNA sequences (Geneious Prime 2019.2.3, https://www.geneious.com). Sequences were run through the NCBI Nucleotide Basic Local Alignment Search Tool (BLASTn) and identified to the genus level, by selecting the top ranked match. Microbial genera were then compared across years, genotypes, and locations, as well as by viable and non-viable seed samples. Figures were also created in R Studio. All packages used in R Studio can be found in Table S3.

### Anti-Microbial Seed Treatments

#### Plant Materials

To evaluate the influence of microbial growth on seed quality in NWR during storage, chemical treatments were utilized in an attempt to reduce microbial growth. Three varieties of NWR, Barron, Itasca-C12, and Itasca-C20, and one elite breeding line, FY-C20, from two locations, Grand Rapids and Clearbrook, MN, tested in 2018 were utilized for this experiment. Before any seed treatments took place, the microbes associated with these genotypes were isolated and reported in the Microbial Profiling section.

#### Antimicrobial Seed Treatments

Two antimicrobial agents, 10% bleach and 10% hydrogen peroxide, were evaluated for two different rinse lengths, 1 min and 5 min. In order to improve the contact angle of the bleach solution, 100 μl of Tween 20, a nonionic surfactant, was added to produce a 10% bleach, 0.01% Tween solution.

Three replications of 100 seeds of each genotype per location were placed in 118 ml, 5.08 cm diameter, polystyrene jars. Prior to antimicrobial treatment, seeds were rinsed in 70% ethanol for 1 min to aid in the breakdown of the seed’s waxy coating, and then tripled rinsed with deionized water. Seeds were then treated with either the 10% bleach solution or 10% hydrogen peroxide for either 1 min or 5 min, and then tripled rinsed with deionized water. For storage, seeds were covered in deionized water and placed in the dark at 3°C for 27 weeks until germination testing. For the control treatment, seeds were pretreated with 70% ethanol, triple-rinsed with deionized water, covered with deionized water, and placed in storage. There were 120 jars in total (4 genotypes x 2 locations x 5 treatments x 3 replications).

#### Post-Treatment Microbial Profiling

The microbes that survived antimicrobial treatments were isolated 32 weeks after treatment. Three replications of three seeds per treatment by location by genotype combination were placed on both PDA and NA. Methods for microbial isolation followed the pre-treatment isolation methods described above.

#### Germination Testing

Viability of treated seed via germination testing was analyzed 27 weeks after treatment. Forty seeds from each treatment jar were placed in a ULINE 2mil 3”x5” plastic sample bag and filled with 10 ml deionized water. Bags were placed under a benchtop grow light programmed for 16-hour days with an ambient temperature of 22°C. Seeds were monitored for 14 days for germination counts, where germination was defined as coleoptile emergence of 1 cm. Germination was also estimated the following year (~90 weeks after treatment) using the seed in storage (Table S2).

#### Data Analysis

Analysis of 16S rRNA and ITS DNA sequences were analyzed with Geneious Prime. Sequences were trimmed and aligned to known fungal and bacterial sequences using the Basic Local Alignment Search Tool (BLAST). Analysis of variance was generated using R (R Core Team, 2017) with fixed effects of location, replication, genotype, and treatment. Using the LSD test function with the Bonferroni adjustment, the least significant differences were calculated for location, genotype, and treatment (Table S3).

## Results and Discussion

### Microbial Profiling of NWR Seed

Host-associated microbiomes are complex communities that can improve our understanding of important biological functions of the host. Research in the area of seed-associated microbiomes has been developing rapidly with a large emphasis on agricultural crops, including *Triticum, Zea*, and *Oryzae* (Johnston-Monje and Raizada, 2011; Eyre et al., 2019; Kuźniar et al., 2020). In this study, we sought to characterize the presence and changes in microbial communities that exist within low temperature NWR seed storage. Bacteria and fungi associated with seeds were isolated from four NWR genotypes grown at three cultivation sites and over a 5-year period. Across all genotypes, locations, and years, 27 total microbial taxa were isolated and identified to the genus level, including 13 fungi and 14 bacteria, which were spread across 5 divisions and 11 classes. We found 13 genera belonging to seven different orders in the Ascomycota (48.1%), one genus in the Mucoromycota (3.7%), two genera from one order in Bacteroidetes (7.4%), two genera from one order in Firmicutes (7.4%), and nine genera spread across five different orders in the Proteobacteria (33.3%) (Table 1). Across samples, the most prevalent Ascomycetes included *Penicillium* (26 samples), *Geotrichum* (15), and *Candida* (9). Proteobacteria including *Pseudomonas* (26) were also prevalent. Only two genera, *Pseudomonas* and *Penicillium*, were found across years, locations, and genotypes (Table S1). Both genera are considered ubiquitous generalists and are commonly isolated from a variety of substrates and locations (Visagie et al., 2014; Chakravarty and Anderson, 2015). An additional 30 isolates were not identified due to difficulty during DNA extractions or anomalies during sequencing, which indicates that future studies are needed and may benefit from high-throughput sequencing methods, which would also provide information about non-culturable microbes.

**Table 1.**
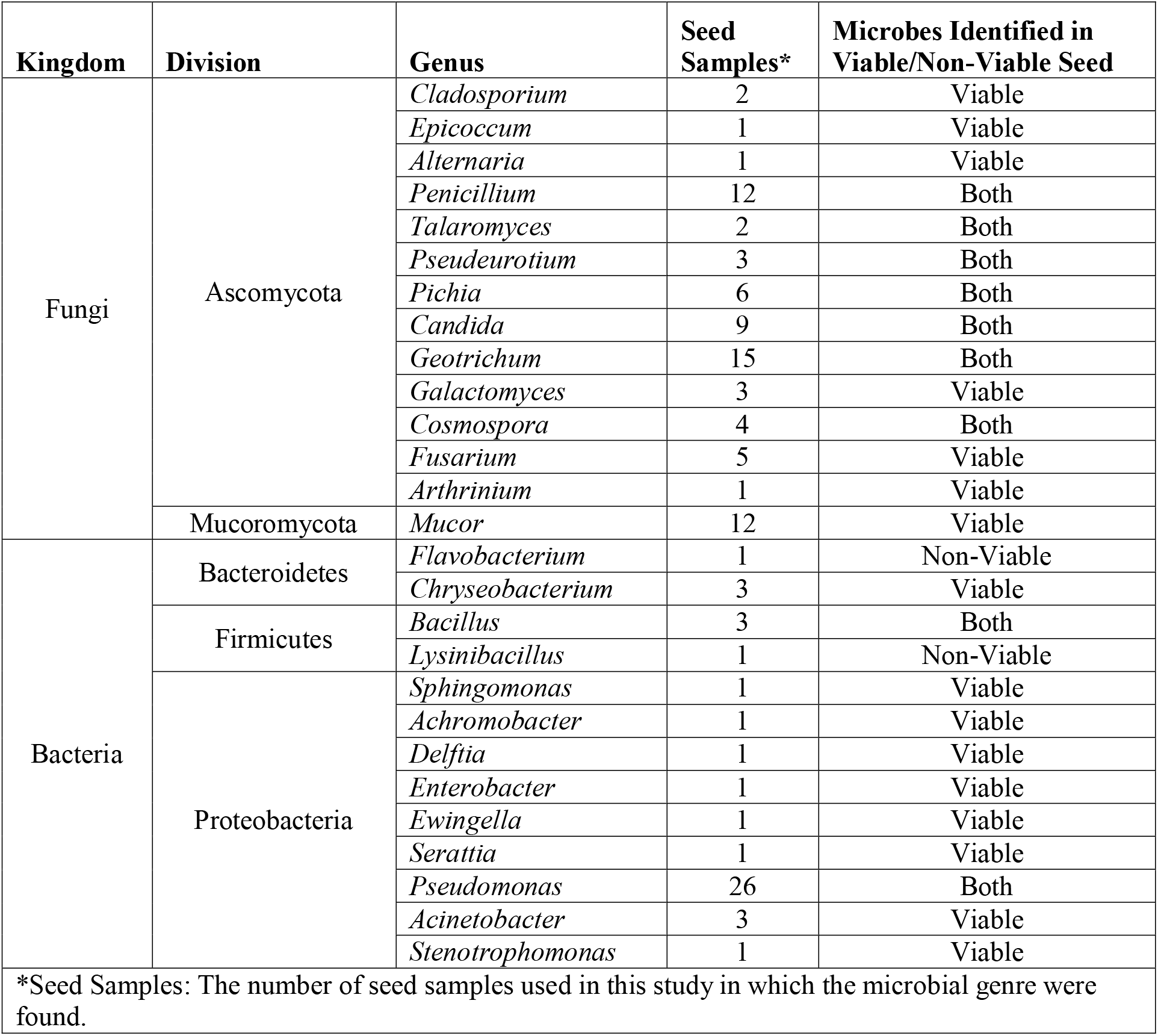
List of fungi and bacteria identified on northern wild rice (NWR; *Z. palustris*) seed stored submerged in water at 3C for 1-4 years. Genera were identified via ITS and 16sRNA sequencing.

### Seed Viability Impacts the NWR Seed Microbiome

The seeds of desiccation-sensitive, recalcitrant species, such as NWR, can only survive in *ex-situ* storage for a limited time (Tweddle et al., 2003; Berjak and Pammenter, 2013). For some species of tropical forest trees, such as *Inga vera* and *Syzygium cumini*, it is only days to months (Vázquez-Yanes and Orozco-Segovia, 1993; Parisi et al., 2016; Nagendra et al., 2019), and for others, like NWR, seed can survive and maintain vigor for ~1 year (Simpson, 1966; Grombacher et al., 1997; Berjak and Pammenter, 2008). During that time, microbial growth on NWR seeds can become abundant (Berjak and Pammenter, 2008). While the presence and consequence of microbes found in orthodox seed storage have been well studied, we know little about the presence and impact of microbial communities in hydrated storage (Calistru et al., 2000). In this study, we found that seed viability played a major role in the microbial constituents of submerged NWR seed. (Table 1; Figure 1a). Due to the short duration of NWR seed viability within our current storage system, we anticipated that seed stored for more than two years would, at best, display low germination. Germination and tetrazolium testing confirmed that all but one sample from 2014 to 2016 seed stocks, representing 3 to 5 years in storage, were non-viable (Figure 1a; Table S2). The average germination rate of seed harvested in 2017, which had been stored for 2 years, was 53.1 % and 33.8 %, respectively, in 2018 (Figure 1a). The average number of microbial genera identified from 2014 to 2016 and 2017 to 2018 seed stocks was 6 and 14, respectively (Figure 1d), demonstrating that the microbial communities from viable seed were more diverse than from non-viable seed. In addition to harboring a more diverse microbiome, 2018 seed also contained the most unique genera, by a large margin (Figure 2a). The microbial genera found in 2017 seed were interesting in that they overlapped with genera found on both viable and non-viable seed from the other years (Figure 2a; Table S1). Therefore, it appears that the microbial community found in 2017 seed, after one year of storage, may represent a transitional phase between viable and non-viable seed. We also identified a shift in the ratio of fungal and bacterial genera, with a roughly 1: 1 ratio in newer seed, but a skew towards more fungal genera in older seed (Figure 1d). These results indicate that microbial diversity in NWR hydrated storage declines with the age, and associated viability of the seed, and that over time, fungi outcompete bacteria. This association between the decline of seed viability and an increase in fungal activity has also been identified in other recalcitrant species (Calistru et al., 2000). The decrease in microbial diversity, following the loss of seed viability, could also be related to a decline in nutrient availability as well as the potential accumulation of antimicrobial substances secreted by the dead organ tissue surrounding the embryo, or produced by other microbes (Godwin et al., 2017; Raviv et al., 2018).

**Figure 1.**
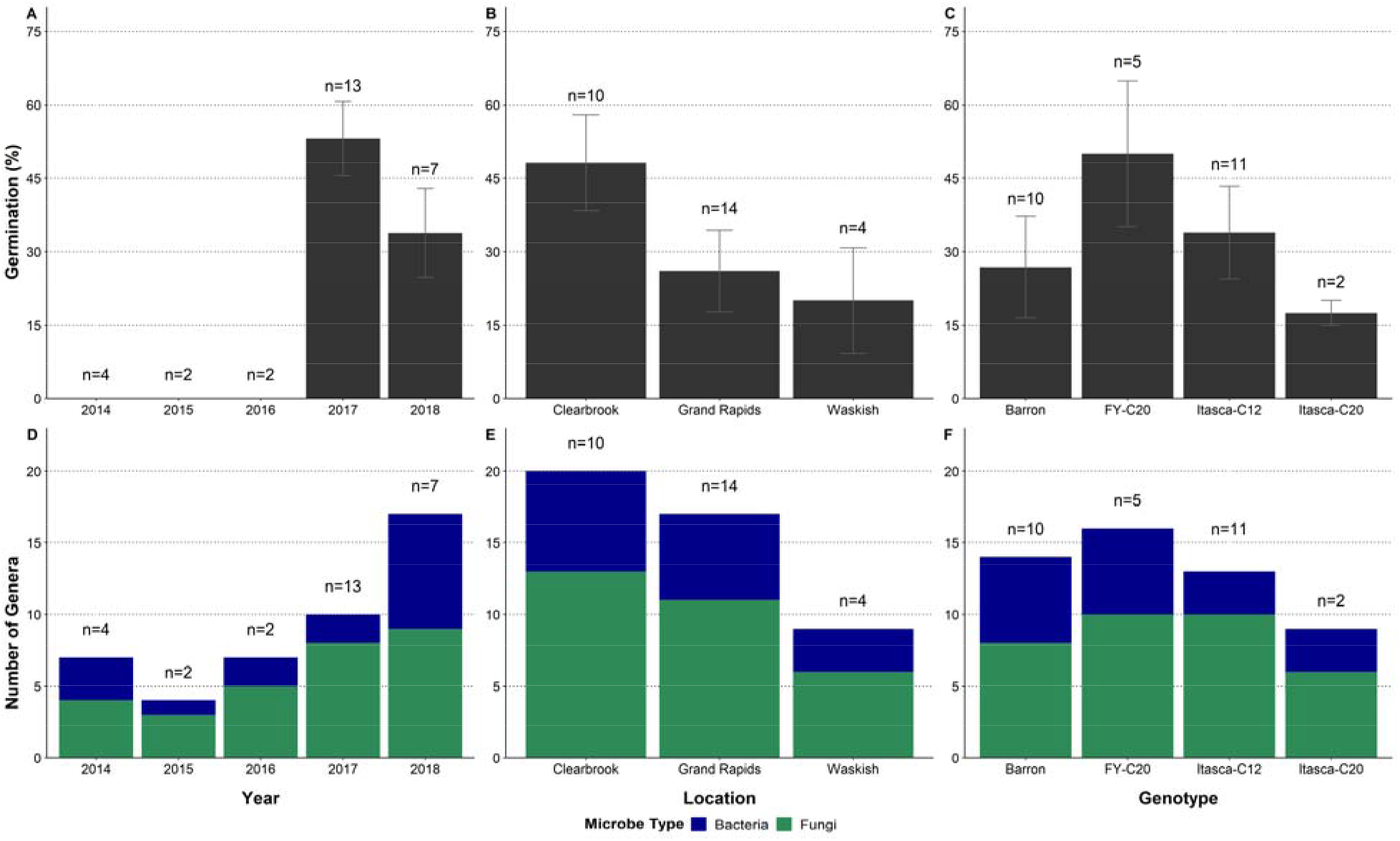
Seed germination rates and number of microbial genera identified on northern wild rice (NWR; *Z. palustris*) seed stored submerged in water at 3°C for 1 to 4 years. Germination rates and number of microbial genera by year of seed harvest (A and C), location of seed production (B and D), and by NWR genotype (C and E) are shown. The number of samples evaluated are indicated above the bars.

### Temporal Analysis of Microbial Communities in NWR Storage

Storage time appears to have a dramatic effect on the composition and diversity of microbial growth on NWR seed. In this study, we assessed the microbial communities from NWR seed stocks that were in storage for 1 to 5 years. Microbes were identified from all seed stocks, regardless of viability, which allowed us to investigate putative shifts in microbial communities throughout storage times. There were nine microbial genera found in both viable and non-viable seeds: the fungal genera *Penicillium*, *Talaromyces*, *Pseudeurotium*, *Pichia*, *Candida*, *Geotrichum*, and *Cosmospora* and the bacterial genera *Bacillus* and *Pseudomonas* (Table 1). Many of these genera are ubiquitous, including *Penicillium*, *Pseudomonas* and *Bacillus* and have been associated with a number of host microbiomes (Links et al., 2014; Visagie et al., 2014; Adam et al., 2018; Eyre et al., 2019). Although found in both viable and non-viable seed, *Talaromyces, Pichia*,and *Geotrichum* species, have been associated with the break-down of seed coats, dehiscence, and decomposition (Telli-Okur et al., 2008; Kim et al., 2017; Sfakianakas et al., 2007). *Flavorbacterium*, is known to efficiently degrade biopolymers and dissolved organic matter, which becomes available as plants decompose (Bernardet and Bowman 2006; Hur et al., 2011).

There was a larger diversity and number of microbes unique to seed from 2017 and 2018, representing the shortest storage lengths, than other years (Figure 1d). A total of 16 genera were unique to either 2017 or 2018 seeds including seven fungal and nine bacterial genera, which were spread out across four divisions and eight classes (Table 1). Of these, only three fungal genera, *Candida, Galactomyces* and *Pseudeurotium*, were unique to 2017 seed. There were four genera found in both 2017 and 2018 seed: *Pseudomonas* and *Penicillium*, which are ubiquitous, as well as *Chryseobacterium* and *Mucor*. Of the unique microbes from 2017 and 2018, some putative functions were identified. The promotion of plant growth has been found for species of *Achromobacter* (Corsini et al., 2018; Jiménez-Vázquez et al., 2020), *Acinetobacter* (Shi et al., 2011), *Chryseobacterium* (Singh and Goel, 2015), *Delftia* (Suchan et al., 2020), and *Stenotrophmonas* (Schmidt et al., 2012). A few of these unique microbes have also been assessed for their production of antimicrobial substances or use as biocontrol agents including species of *Delftia* (Prasannakumar et al., 2015), *Enterobacter* (Gong et al., 2019), *Serratia* (Johnson et al., 2001), and *Sphingomonas* (Wachowska et al., 2013). Several of the fungal genera identified here, including species of *Alternaria* (Casa et al., 2012; Gambacorta et al., 2018), *Cladosporium* (Schenck and Stotzky, 1975), *Epicoccum* (Schenck and Stotzky, 1975; Anžlovar et al., 2017), *Fusarium* (Mycock and Berjak, 1990; Gambacorta et al., 2018), and *Penicillium* (Mycock and Berjak, 1990; Gambacorta et al., 2018) commonly accumulate during grain seed storage, with some species ultimately leading to seed deterioration or the production of mycotoxins (Magan et al., 2004; Links et al., 2014). Still others are pathogenic to a variety of hosts including species of *Alternaria* (Maude and Humpherson-Jones, 1980; Babadoost, 2011), *Cladosporium* (Kirk and Crompton, 1984; Nam et al., 2015), *Epicoccum* (Stokholm et al., 2016), *Ewingella* (González et al., 2012), *Fusarium* (Babadoost, 2011), and *Galactomyces* (Song et al., 2020). Ultimately, due to the complexity of host-microbe interactions, it is difficult to draw broad conclusions about the function of these microbes in NWR seed storage. Further testing of microbial interactions in NWR is needed to clarify the true function(s) of these microbes in this unique ecosystem.

### Spatial Analysis: Microbes across Locations and Genotypes

Environmental factors, such as local climate, soil composition, nutrient availability, and water, which vary based on growing region or even field location, all can affect the microbial community composition associated with plants (Buyer, 1999; Hacquard, 2016). Within this study, we evaluated three different sites across northern Minnesota common to NWR cultivation and identified differences in the number of genera among locations (Figure 2b). Five genera were common to all locations, possibly indicating that these microbes are found within, rather than on the surface of seed, and could be part of a core NWR microbiome. These included *Bacillus*, *Candida*, *Geotrichum*, *Pichia*, and *Pseudomonas*. Many of these microbes have been associated with the deterioration or decomposition of various tissues in other species, such as Orchidea (Sfakianakis et al., 2007), several fruits species, (Talibi et al., 2012; Nishikawa et al., 2006), and sunflower (Telli-Okur and Eken-Saracoglu, 2008). The low number of unique microbes identified in Waskish could be due to the lower number of samples tested (n=4, compared with Clearbrook, n=10, and Grand Rapids, n=14) (Figure 1a). Each location had their own management practices as well as unique weather conditions, which may have influenced the microbes identified at these locations. While location-specific differences in microbial communities are important to understand, they are just one piece in the complex interaction between environmental conditions and host species, and their influence on microbial communities.

**Figure 2.**
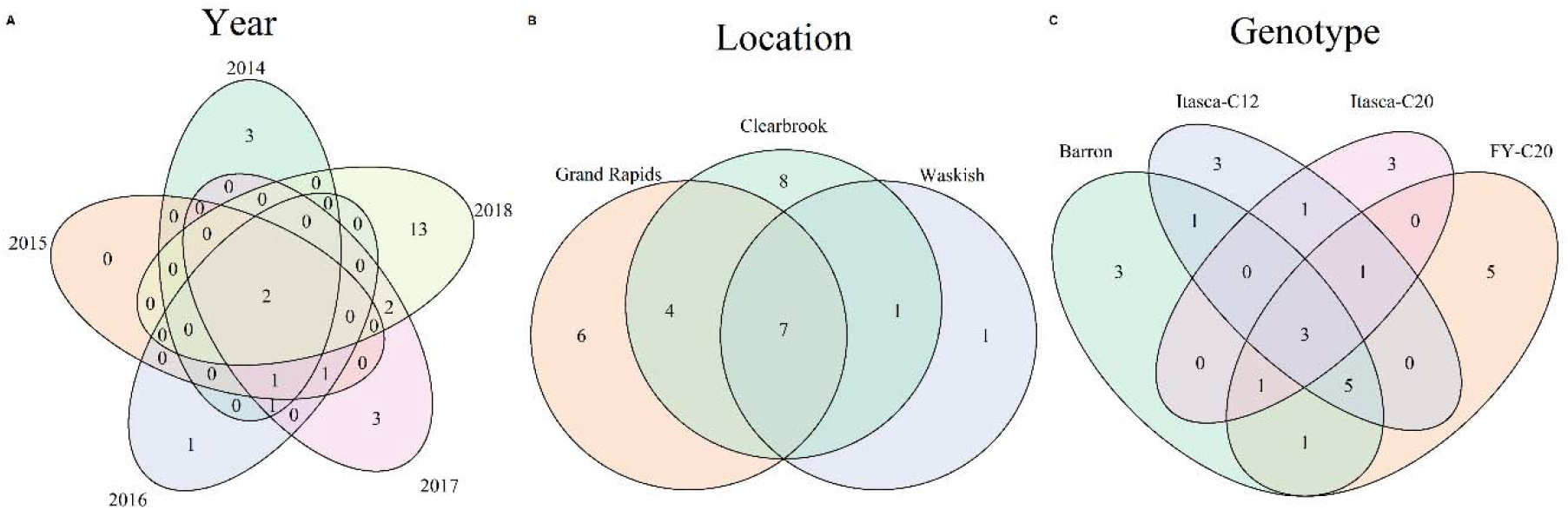
Venn diagrams depicting shared and unique microbes identified on northern wild rice (NWR; *Z. palustris*) seed stored submerged in water at 3°C for 1 to 4 years by A) year, B) location, and C) genotype.

Some plant microbial communities are vertically transmitted from generation to generation, which can lead to genotype-specific differences in microbial communities (Johnston-Monje, 2016; Adam, 2016; Nelson, 2004; Shade, 2017). We evaluated four different genotypes in this study to identify potential differences between genotypes of NWR. Similar numbers of microbial genera were found in Barron, FY-C20, and Itasca-C12 samples (14, 16, and 14, respectively) (Figure 1e). By comparison, only nine genera were isolated from Itasca-C20 samples. The lower diversity of microbes in Itasca-C20 could be due to the low sample number from this variety that was used in this study (n=2). *Mucor*, as well as *Penicillium* and *Pseudomonas*, were found in all genotypes. *Acinetobacter*, *Bacillus*, *Candida*, *Chryseobacterium*, *Cosmospora*, *Geotrichum*, and *Pichia* were found in three out of the four genotypes. We did not identify any consistent relationships in microbial communities based on NWR genotype when exploring the interaction between genotype, location, and year. The microbial communities seem to be largely dictated by viability of seed and more testing on viable seed, in general, is needed to confirm these findings.

### Antimicrobial Seed Treatments Reveal Microbial Communities in NWR Seeds Can Shift Rapidly

This study sought to identify the effect of seed treatments on seed germination and diversity of microbes present on NWR seed following storage. For this experiment, the microbes identified from the 2018 samples, described in the previous section, were compared to microbial communities isolated from untreated seed and seed treated with antimicrobial solutions. Following treatment and 32 weeks in storage, seed germination and microbial growth were assessed for each treatment, genotype, and location. Significant differences in germination among locations, genotypes, and the interactions of treatment x location and location x genotype were found in this study (Table 2). Although the ANOVA indicated that there was a significant interaction between treatment and location, the only differences found using a post-hoc LSD test were due primarily to differences by location. This implies that any differences resulting from treatments were quite small.

**Table 2.** Analysis of Variance (ANOVA) for the assessment of the influence of five antimicrobial seed treatments on the germination of four seed samples of northern wild rice (NWR; *Z. palustris*) from two locations in 2018.

|  | Df | Sum Sq | Mean Sq | F value | Pr(>F) |
| --- | --- | --- | --- | --- | --- |
| Reps | 2 | 459.36 | 229.68 | 2.39 | 0.0988 |
| Treatments | 4 | 322.08 | 80.52 | 0.84 | 0.5063 |
| Locations | 1 | 16635.10 | 16635.10 | 172.76 | <0.0001 |
| Genotypes | 3 | 15219.84 | 5073.28 | 52.69 | <0.0001 |
| Treatment x Location | 4 | 1056.86 | 264.22 | 2.74 | 0.0342 |
| Treatment x Genotype | 12 | 1245.62 | 103.80 | 1.08 | 0.3901 |
| Location x Genotype | 3 | 5974.55 | 1991.52 | 20.68 | <0.0001 |
| Trt. x Loc. x Genotype | 12 | 1762.13 | 146.84 | 1.52 | 0.1331 |
| Residuals | 78 | 7510.76 | 96.29 | N/A | N/A |

The microbiome from untreated seed differed from that of the control and treated seed after a 32 week storage period (Table 3). Seven microbial genera were found only in the untreated seed: *Achromobacter*, *Arthrinium*, *Delftia*, *Enterobacter*, *Epiccocum*, *Ewingella*, and *Penicillium*. Because these genera were not found in control seed, their loss potentially represents a natural shift in microbial diversity with submerged storage. Additionally, there were 13 microbial genera found in the control but not untreated seed. Ultimately, untreated seed and control seed shared only five of 31 identified microbial genera: *Chryseobacterium*, *Fusarium*, *Pseudomonas*, *Serratia*, and *Sphingomonas* (Table 3). This dramatic shift in microbial community composition, between untreated and control seed, may reflect the change from a dry environment to submerged storage, which decreases the availability of oxygen, changes in nutrient availability, or the effect of seed germination, following the natural dormancy period, and the associated exudates (Nelson, 2004; Barret et al., 2015; Truyens et al., 2015). However, to the best of our knowledge, there are no published studies comparing the microbiome of recalcitrant seed, before and following submerged storage that would aid in the development of more concrete conclusions about the observed phenomenon.

**Table 3.**
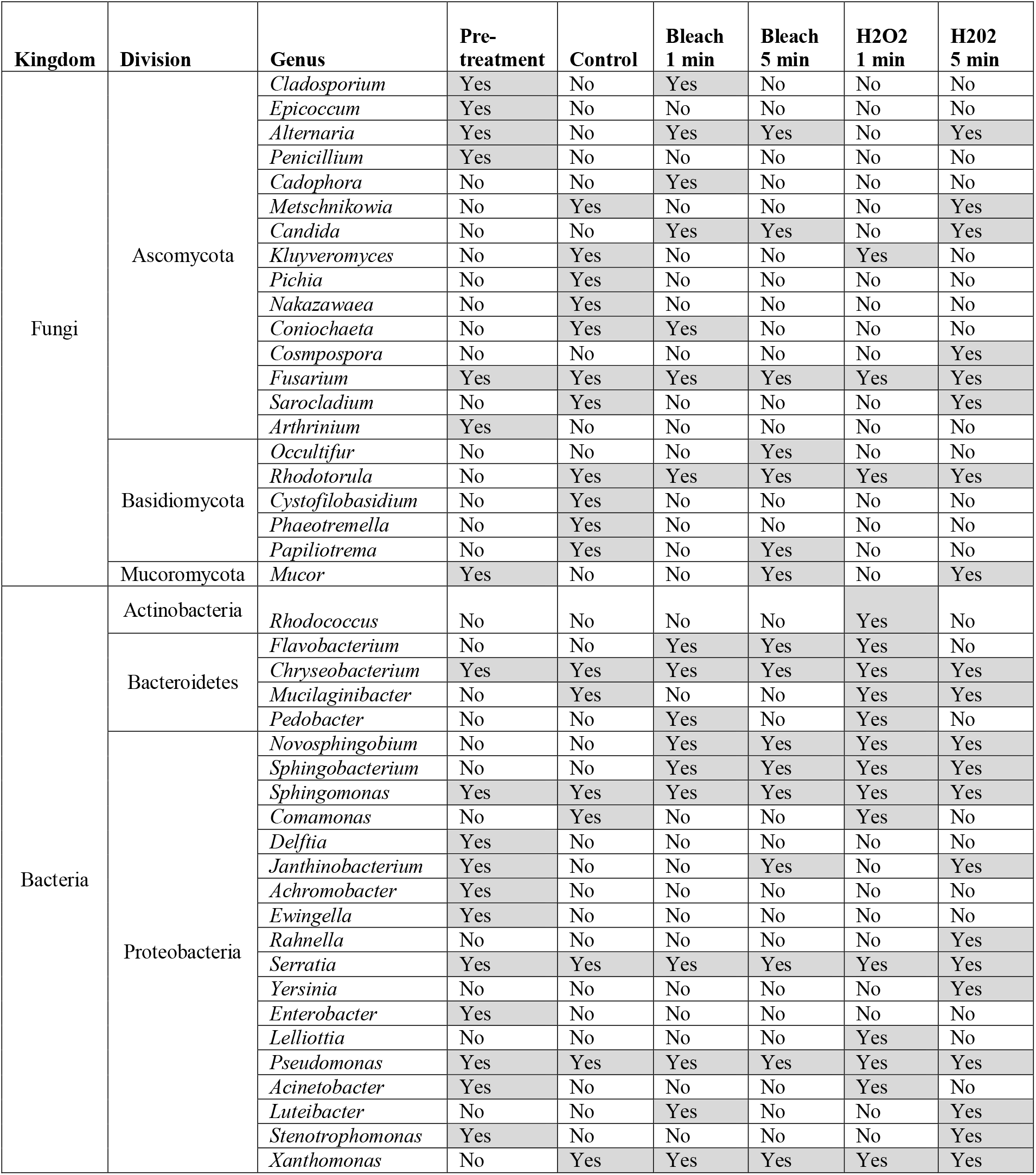
List of microbes identified post-application of antimicrobial seed treatments on northern wild rice (NWR; *Z. palustris*) seed. Seed was collected in 2018 and stored submerged in water at 3°C for ~8 months post-treatment before testing for germination. Yes indicates that the genus was found on seed within the defined treatment and No that it was not found within that treatment.

### Antimicrobial Seed Treatments Did Not Have A Strong Effect on Germination or Microbial Growth

The seed treatments used in this study did not appear to strongly influence seed germination or the microbes associated with seed, when compared to the control. There were no statistical differences in the germination of control versus treated seed or between seed treatments (Table 2). Likewise, it appears that the loss of microbial genera, compared to control seed, cannot be linked to any of the tested treatments, as there were no microbial genera shared by the control and untreated seed that were not also found in the treated seed. However, 13 microbial genera were found in treated seed but not in the control, suggesting that treatments allowed for the growth of otherwise undetected microbes. Surface sterilization via antimicrobial treatments may effectively control surface microbes, which may also create favorable conditions for internal microbes to occupy external seed tissues (Blanchard, 1973; Mohamed, 2017).

While similar numbers of microbial genera were present across all treatments (16 to 21), only seven of these genera were shared with genera from the control: *Chryseobacterium*, *Fusarium*, *Pseudomonas*, *Rhodotorula*, *Serratia*, *Sphingomonas*, and *Xanthomonas*. These microbes represent a core microbiome that was unaffected by the seed treatments, which matches closely with the core microbes identified between untreated and control seed. *Pseudomonas*, *Sphingomonas*, and *Fusarium* are genera commonly found in the core microbiome of crop species including *Cucurbita pepo*, *Brassica napus*, *Oryza sativa*, *Triticum aestivum*, and *Lens culinaris* (Rybakova et al., 2017; Adam et al., 2018; Eyre et al., 2019; Morales Moreira et al., 2020). It has been noted previously that *Pseudomonas* and *Fusarium* are not always eliminated from seed following treatment with sodium hypochlorite (Cuero et al., 1986). However, other seed treatments have been efficacious in the elimination or reduction of fungi commonly associated with recalcitrant seed, including *Penicillium* sp., *Cladosporium* sp., and *Fusarium* sp. (Oliveira et al., 2011; Françoso and Barbedo, 2014; Parisi et al., 2016). This suggests that different seed treatments than those tested here may be more effective in the reduction of microbial growth during submerged NWR seed storage.

### The Effect of Environment and Genotype on the Microbial Constituents Post-Antimicrobial Treatments

Location appeared to influence the germination and microbial constituents of seed in this study. Averaged across treatments, Clearbrook seed had significantly higher germination than Grand Rapids (Figure 3a). This may be related to the health of the parental plants, which were grown under differing environmental conditions and management practices. Control seed was used to identify the effect of location on the microbial diversity of seed. Only 3 of 17 microbial genera were found in seed from both locations (Table S4). This implies that the parental environment may have influenced the seed microbiomes. Previous research has found differences in microbial diversity of crops including *Phaseolus vulgaris*, *Triticum aestivum*, *Brassica napus*, *Lens culinaris* seed between locations, owing both to differences in the growing environment as well as the management of the land (Barret et al., 2015; Hardoim et al., 2015; Klaedtke et al., 2016; Morales Moreira et al., 2020).

**Figure 3.**
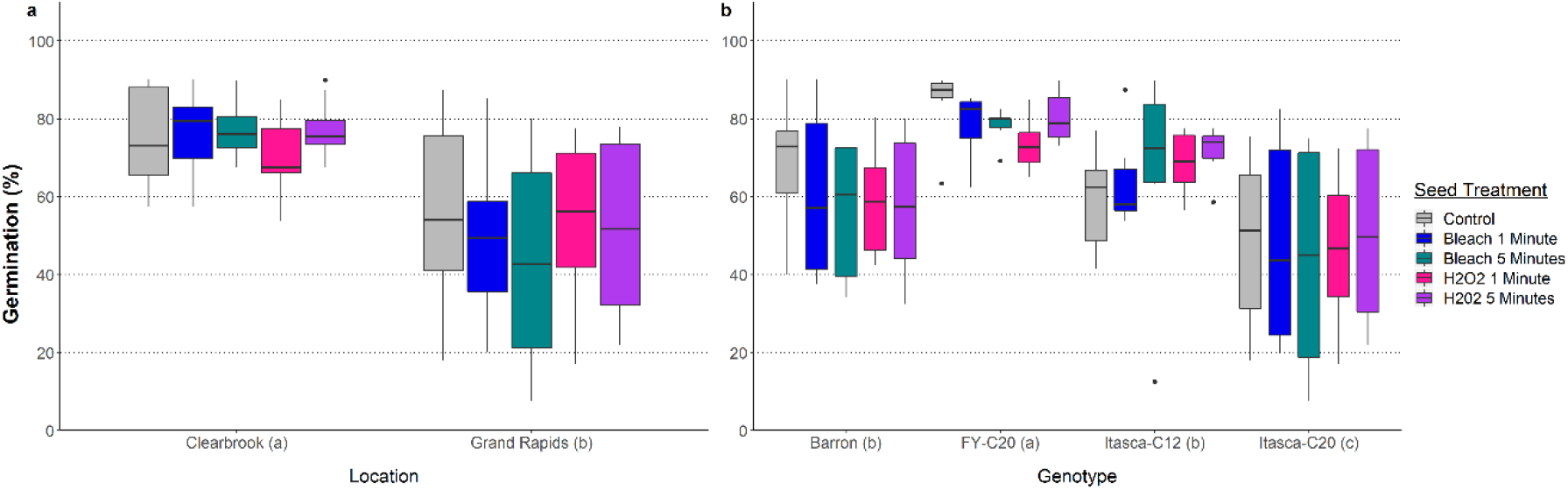
The germination rates of seed from four northern wild rice (NWR; *Z. palustris*) genotypes from two locations after ~8 months in hydrated storage at 3°C and post-application of antimicrobial seed treatments. The letters next to the x-axis labels indicate significant differences that were present between locations and genotypes.

**Figure 4.**
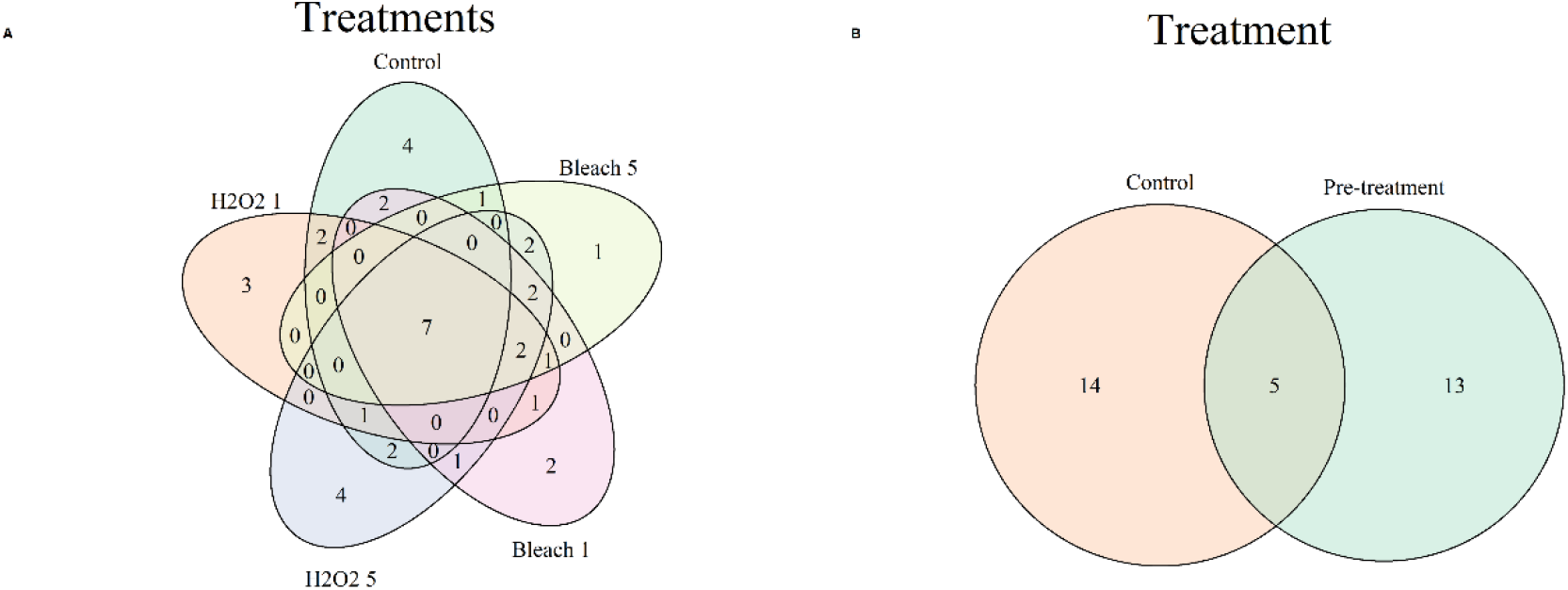
Venn diagrams depicting shared and unique microbes identified on northern wild rice (NWR; *Z. palustris*) seed stored submerged in water at 3°C for ~ 8 months post-application of antimicrobial seed treatments. A) Comparison of control and treatments. B) Comparison of untreated vs. control.

Genotype affected both the germination and the microbial communities present on seed. When averaged across treatments, FY-C20 had the highest germination, followed by Barron and Itasca-C12, and finally, Itasca-C20 (Figure 3b). Although differences were found between genotypes, it should be noted that the germination of NWR varieties often fluctuates between years. Therefore, it is possible that across years, these differences would be decreased or eliminated. The control seed lots were used to compare the natural progression of storage microbiota over time, between the four NWR genotypes. Only two of 17 genera were shared between all genotypes: *Pseudomonas* and *Sphingomonas*, with an additional four genera shared between at least two of the genotypes: *Chryseobacterium, Fusarium, Pichia*, and *Serratia*. Together, these shared genera make up the core microbiome as described above. This indicates that the core microbiome remains mostly unchanged between genotypes but that genotypes do harbor different microbes. It is not uncommon for genotype to play a role in the microbiome of seed, in some cases due to the transmission of endophytes by the parental plant, as has been seen in other plant species (Morales Moreira et al., 2020; Shade et al., 2017; Tannenbaum et al., 2020).

### Conclusions

In this study, we investigated the culturable fungi and bacteria associated with NWR hydrated seed storage over time and space. Microbial constituents were largely dependent on the viability of NWR seed with more genera associated with viable than nonviable seeds. The number of genera declined over longer periods of storage, shifting towards fungal-dominant communities. With the molecular tools utilized in this study it was not possible to identify microbes to the species level, therefore it is unclear whether isolates may have been pathogenic. However, *Pseudomonas syringae* and multiple species in the genus *Fusarium* are known to cause bacterial leaf streak and Fusarium head blight of wild rice, respectively (Bowden and Percich, 1983; Nyvall et al., 1999). Therefore, it may be useful to further characterize the *Fusarium* and *Pseudomonas* isolates that were found in this study. The antimicrobial treatments analyzed were not effective in reducing the number of unique microbes present, which may support the hypothesis that microbes within the seed endosperm and embryo are opportunistic and will occupy the external seed tissues. Further analysis into the timing of treatment in addition to the type of treatment is necessary. It is also unclear at this time whether microbial growth led to a decrease in germination of NWR seed or the reduction of NWR seed viability increased microbial growth, and further testing is needed. Ultimately, this study highlights the difficulty of controlling microbial growth in storage conditions for aquatic, recalcitrant species and improved storage methods may be necessary to effectively control their growth.

## Acknowledgements

The authors would like to thank Deborah Samac for the careful review of this manuscript.

## SUPPLEMENTAL TABLES

**Table S1.**
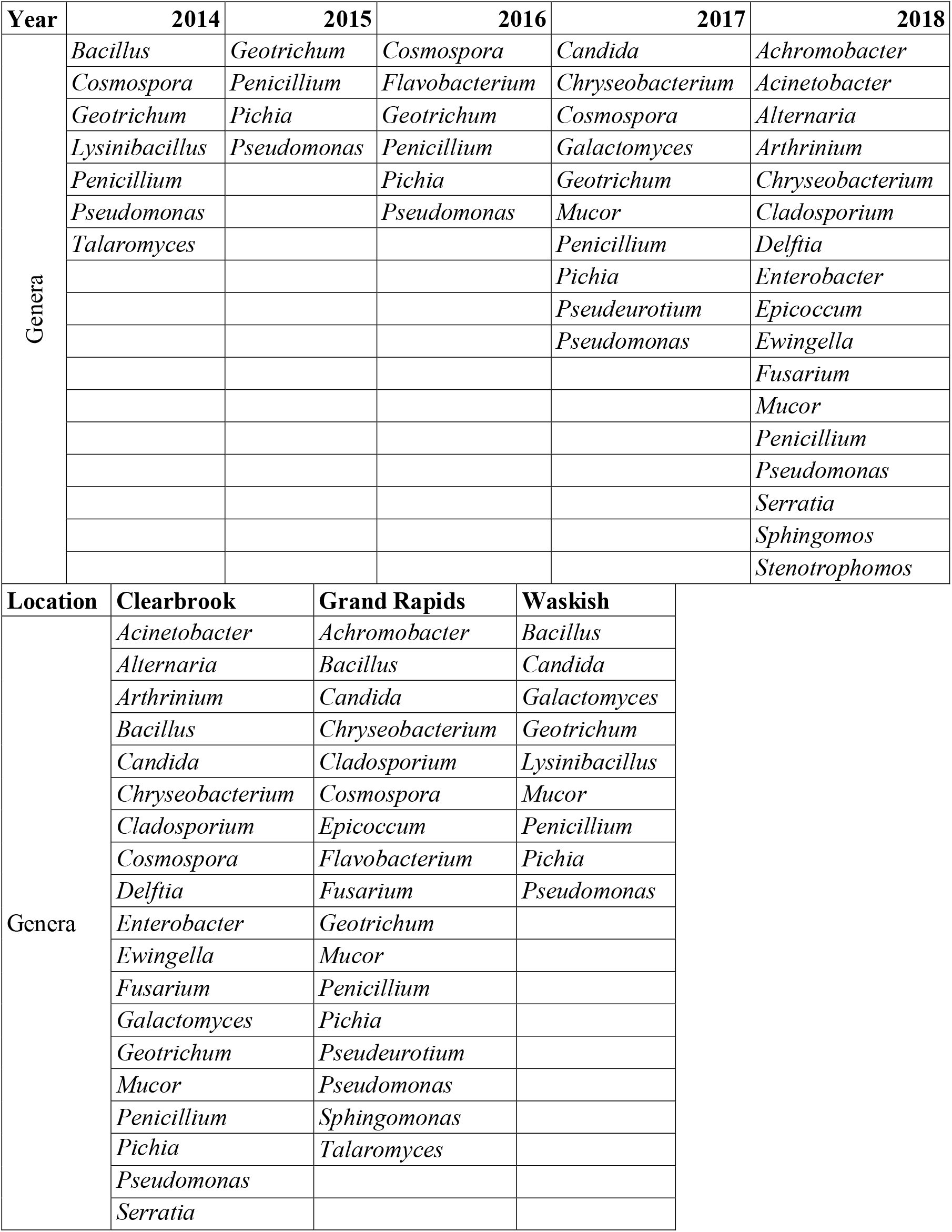
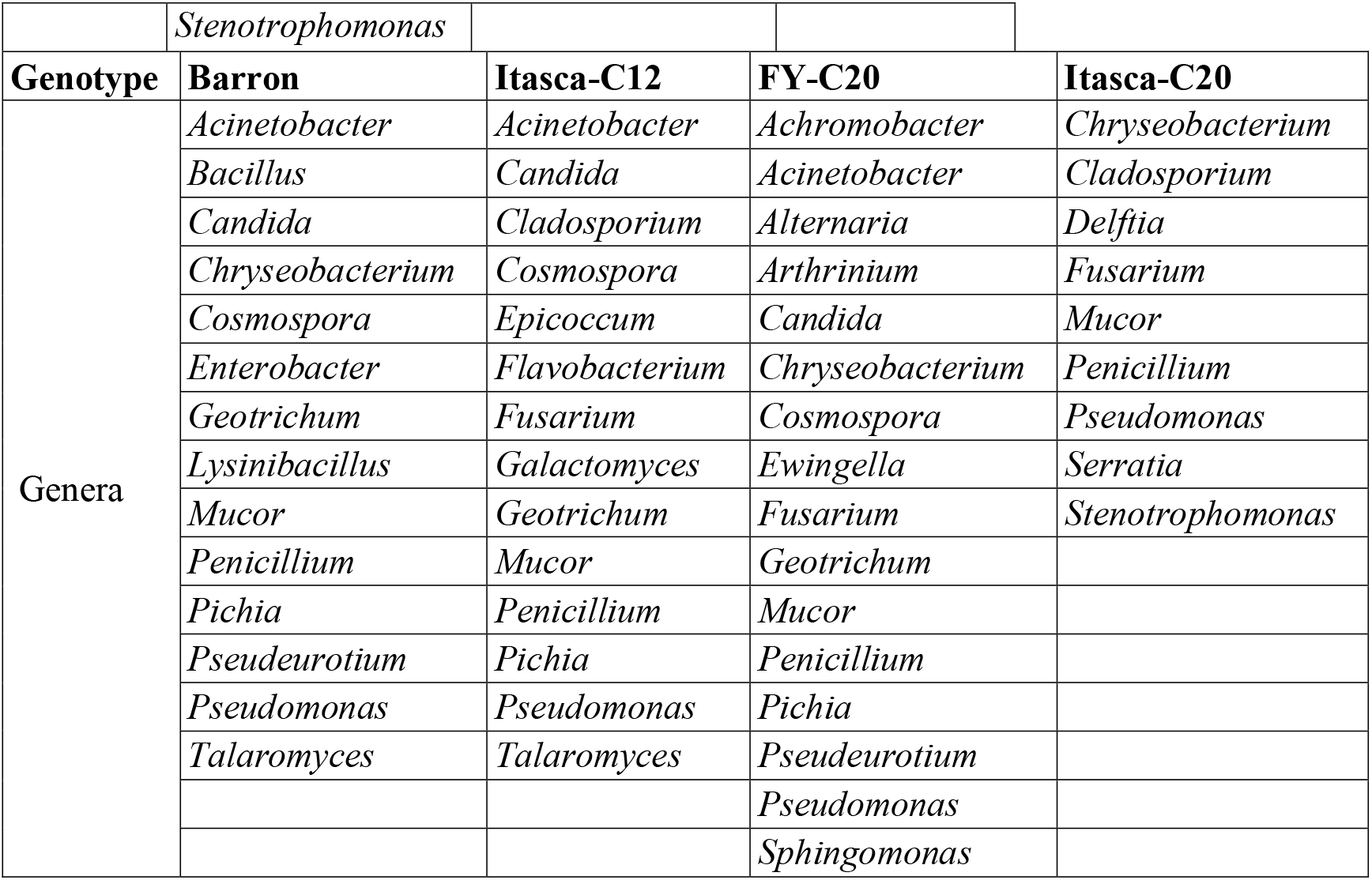
Lists of microbes identified on northern wild rice (NWR; *Z. palustris*) seed stored submerged in water at 3°C for 1-4 years. Genera were identified via ITS and 16sRNA sequencing.

| Year | 2014 | 2015 | 2016 | 2017 | 2018 |
| --- | --- | --- | --- | --- | --- |
| Genera | *Bacillus* | *Geotrichum* | *Cosmospora* | *Candida* | *Achromobacter* |
|  | *Cosmospora* | *Penicillium* | *Flavobacterium* | *Chryseobacterium* | *Acinetobacter* |
|  | *Geotrichum* | *Pichia* | *Geotrichum* | *Cosmospora* | *Alternaria* |
|  | *Lysinibacillus* | *Pseudomonas* | *Penicillium* | *Galactomyces* | *Arthrinium* |
|  | *Penicillium* |  | *Pichia* | *Geotrichum* | *Chryseobacterium* |
|  | *Pseudomonas* |  | *Pseudomonas* | *Mucor* | *Cladosporium* |
|  | *Talaromyces* |  |  | *Penicillium* | *Delftia* |
|  |  |  |  | *Pichia* | *Enterobacter* |
|  |  |  |  | *Pseudeurotium* | *Epicoccum* |
|  |  |  |  | *Pseudomonas* | *Ewingella* |
|  |  |  |  |  | *Fusarium* |
|  |  |  |  |  | *Mucor* |
|  |  |  |  |  | *Penicillium* |
|  |  |  |  |  | *Pseudomonas* |
|  |  |  |  |  | *Serratia* |
|  |  |  |  |  | *Sphingomonas* |
|  |  |  |  |  | *Stenotrophomonas* |
| Location | Clearbrook | Grand Rapids | Waskish |  |  |
| Genera | *Acinetobacter* | *Achromobacter* | *Bacillus* |  |  |
|  | *Alternaria* | *Bacillus* | *Candida* |  |  |
|  | *Arthrinium* | *Candida* | *Galactomyces* |  |  |
|  | *Bacillus* | *Chryseobacterium* | *Geotrichum* |  |  |
|  | *Candida* | *Cladosporium* | *Lysinibacillus* |  |  |
|  | *Chryseobacterium* | *Cosmospora* | *Mucor* |  |  |
|  | *Cladosporium* | *Epicoccum* | *Penicillium* |  |  |
|  | *Cosmospora* | *Flavobacterium* | *Pichia* |  |  |
|  | *Delftia* | *Fusarium* | *Pseudomonas* |  |  |
|  | *Enterobacter* | *Geotrichum* |  |  |  |
|  | *Ewingella* | *Mucor* |  |  |  |
|  | *Fusarium* | *Penicillium* |  |  |  |
|  | *Galactomyces* | *Pichia* |  |  |  |
|  | *Geotrichum* | *Pseudeurotium* |  |  |  |
|  | *Mucor* | *Pseudomonas* |  |  |  |
|  | *Penicillium* | *Sphingomonas* |  |  |  |
|  | *Pichia* | *Talaromyces* |  |  |  |
|  | *Pseudomonas* |  |  |  |  |
|  | *Serratia* |  |  |  |  |

|  |  |  |  |  |
| --- | --- | --- | --- | --- |
|  | *Stenotrophomonas* |  |  |  |
| **Genotype** | **Barron** | **Itasca-C12** | **FY-C20** | **Itasca-C20** |
| Genera | *Acinetobacter* | *Acinetobacter* | *Achromobacter* | *Chryseobacterium* |
|  | *Bacillus* | *Candida* | *Acinetobacter* | *Cladosporium* |
|  | *Candida* | *Cladosporium* | *Alternaria* | *Delftia* |
|  | *Chryseobacterium* | *Cosmospora* | *Arthrinium* | *Fusarium* |
|  | *Cosmospora* | *Epicoccum* | *Candida* | *Mucor* |
|  | *Enterobacter* | *Flavobacterium* | *Chryseobacterium* | *Penicillium* |
|  | *Geotrichum* | *Fusarium* | *Cosmospora* | *Pseudomonas* |
|  | *Lysinibacillus* | *Galactomyces* | *Ewingella* | *Serratia* |
|  | *Mucor* | *Geotrichum* | *Fusarium* | *Stenotrophomonas* |
|  | *Penicillium* | *Mucor* | *Geotrichum* |  |
|  | *Pichia* | *Penicillium* | *Mucor* |  |
|  | *Pseudeurotium* | *Pichia* | *Penicillium* |  |
|  | *Pseudomonas* | *Pseudomonas* | *Pichia* |  |
|  | *Talaromyces* | *Talaromyces* | *Pseudeurotium* |  |
|  |  |  | *Pseudomonas* |  |
|  |  |  | *Sphingomonas* |  |

**Table S2.** List of germination rates of seed from northern wild rice (NWR; *Z. palustris*) genotypes utilized in this study.

| Genotype | Harvest Year | Location | Germination (%) | Metabolically Active* |
| --- | --- | --- | --- | --- |
| Barron | 2014 | Clearbrook | 0 | No |
| Barron | 2014 | Waskish | 0 | No |
| Itasca-C12 | 2014 | Grand Rapids | 0 | No |
| Barron | 2014 | Grand Rapids | 0 | Yes |
| Itasca-C12 | 2015 | Grand Rapids | 0 | No |
| Barron | 2015 | Clearbrook | 0 | No |
| Barron | 2016 | Grand Rapids | 0 | No |
| Itasca-C12 | 2016 | Grand Rapids | 0 | No |
| Barron | 2017 | Grand Rapids | 40 | Yes |
| Itasca-C12 | 2017 | Grand Rapids | 35 | Yes |
| Itasca-C12 | 2017 | Clearbrook | 70 | Yes |
| Itasca-C12 | 2017 | Clearbrook | 65 | Yes |
| Itasca-C12 | 2017 | Clearbrook | 75 | Yes |
| FY-C20 | 2017 | Grand Rapids | 80 | Yes |
| Itasca-C12 | 2017 | Waskish | 10 | Yes |
| Itasca-C12 | 2017 | Waskish | 50 | Yes |
| Barron | 2017 | Waskish | 20 | Yes |
| Barron | 2017 | Clearbrook | 80 | Yes |
| FY-C20 | 2017 | Grand Rapids | 10 | Yes |
| Barron | 2017 | Grand Rapids | 75 | Yes |
| FY-C20 | 2017 | Grand Rapids | 80 | Yes |
| Itasca-C12 | 2018 | Grand Rapids | 5 | Yes |
| FY-C20 | 2018 | Grand Rapids | 20 | Yes |
| Itasca-C20 | 2018 | Grand Rapids | 20 | Yes |
| Barron | 2018 | Clearbrook | 53 | Yes |
| Itasca-C20 | 2018 | Clearbrook | 15 | Yes |
| Itasca-C12 | 2018 | Clearbrook | 63 | Yes |
| FY-C20 | 2018 | Clearbrook | 60 | Yes |

\*Metabolic activity determined by the results of germination and tetrazolium testing

**Table S3.** List of R Studio packages used for analysis and figure creation in this study.

| Package | Documentation |
| --- | --- |
| VennDiagram | https://cran.r-project.org/web/packages/VennDiagram/VennDiagram.pdf |
| png | https://cran.r-project.org/web/packages/png/png.pdf |
| ggpubr | https://cran.r-project.org/web/packages/ggpubr/ggpubr.pdf |
| ggplot2 | https://cran.r-project.org/web/packages/ggplot2/ggplot2.pdf |
| plyr | https://cran.r-project.org/web/packages/plyr/plyr.pdf |
| dplyr | https://cran.r-project.org/web/packages/dplyr/dplyr.pdf |
| agricolae | https://cran.r-project.org/web/packages/agricolae/agricolae.pdf |

**Table S4.**
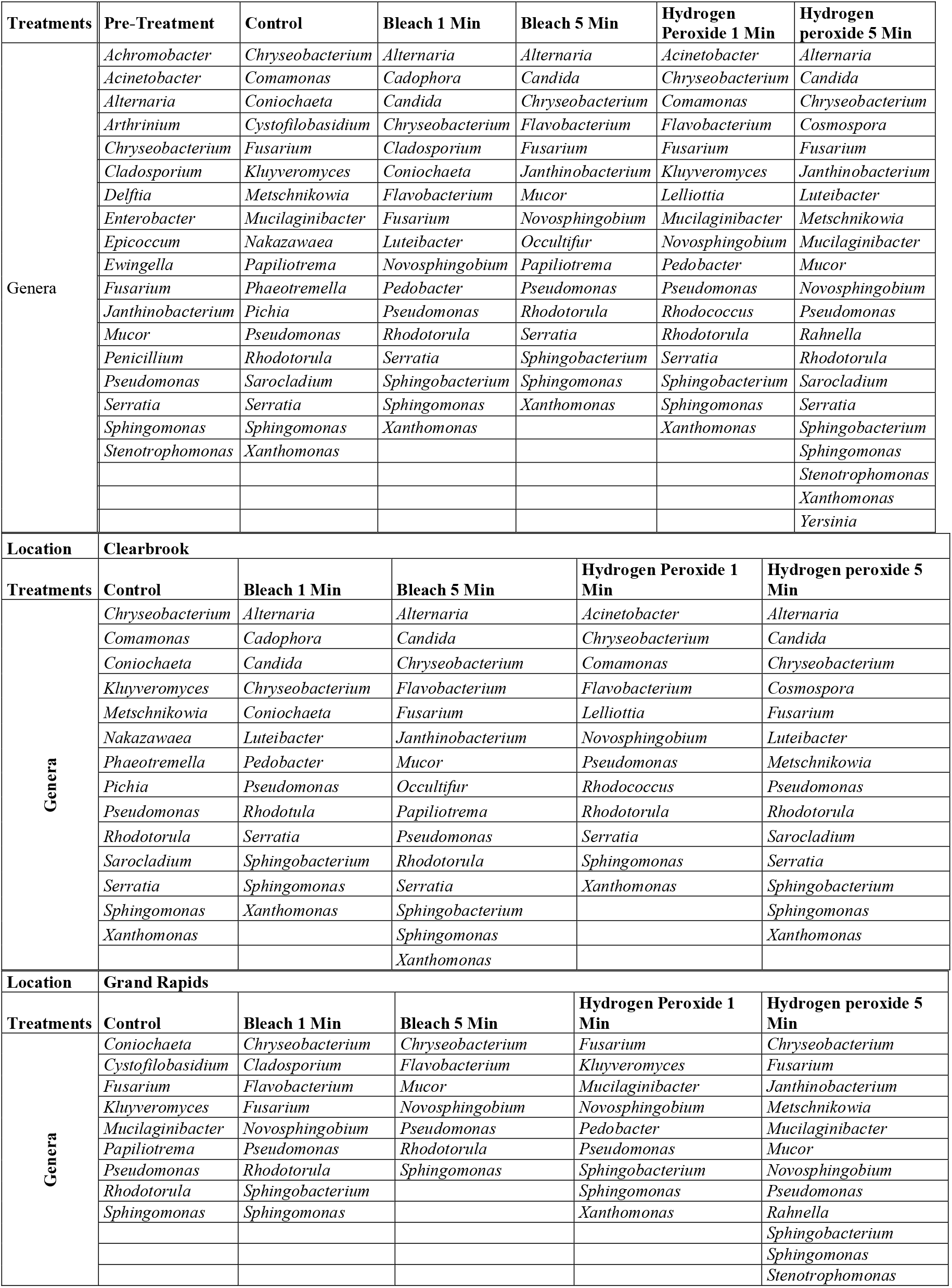
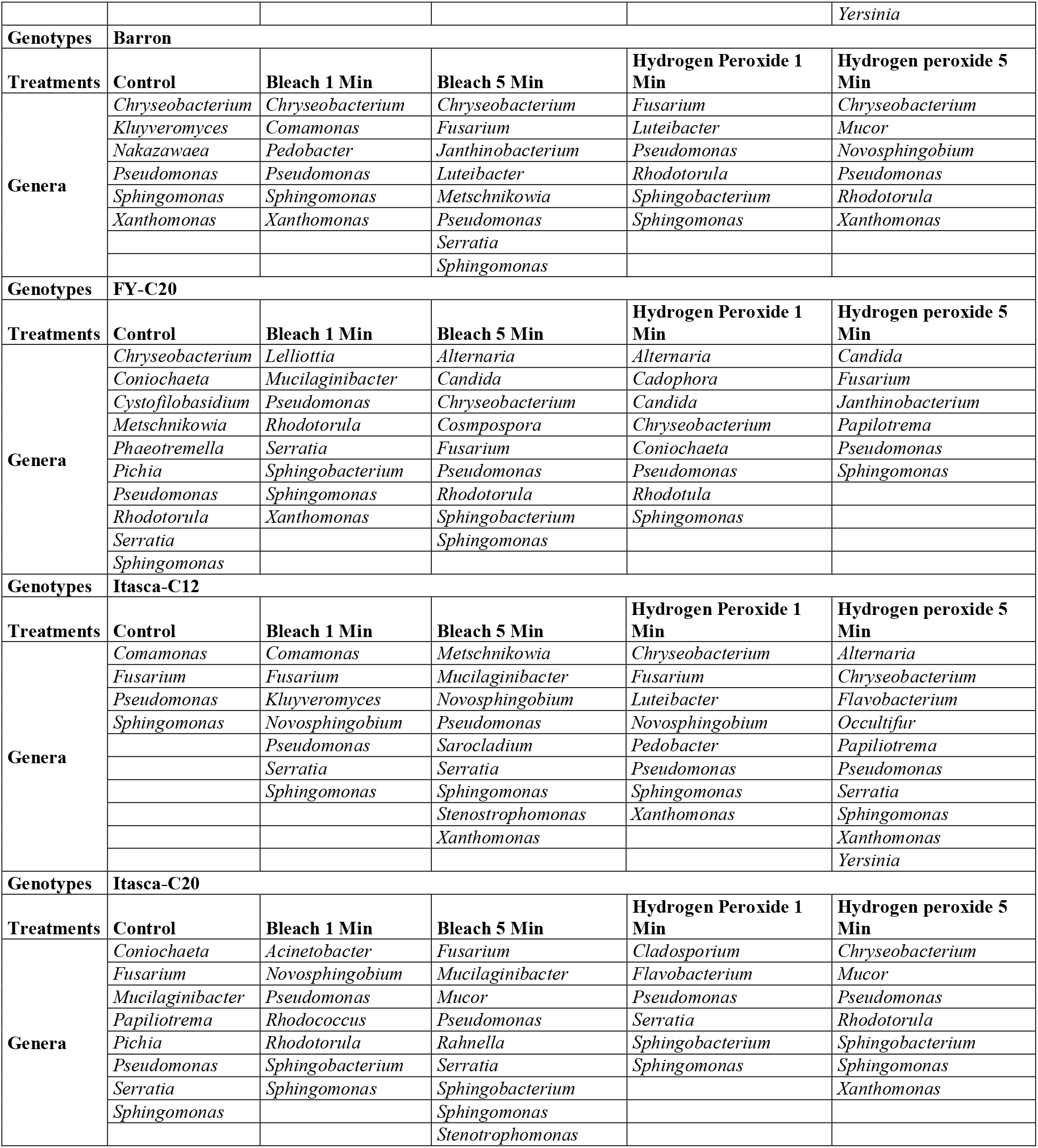
Microbes identified on northern wild rice (NWR; *Z. palustris*) seed stored submerged in water at 3°C post-antimicrobial treatments. Genera were identified via ITS and 16S RNA sequencing.

**Table S5.**
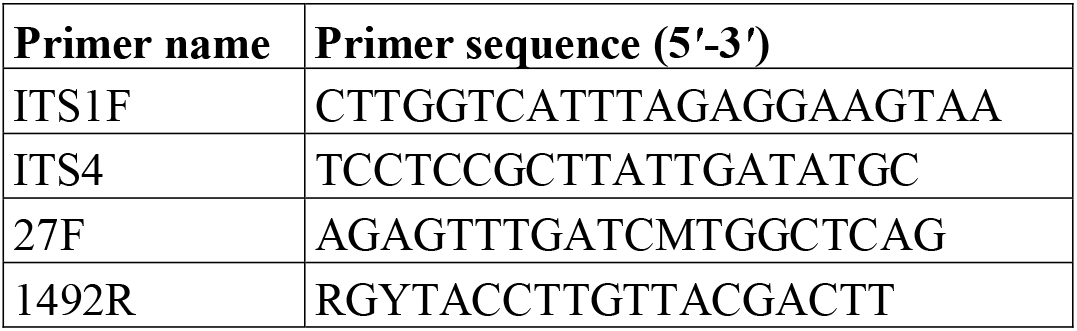
List of ITS and 16sRNA primer sequences used in this study.

